# Technical Note: Focused ultrasound-mediated blood-brain barrier opening for delivery of LNP-packaged modRNA therapy in a mouse model of Niemann-Pick Disease Type C

**DOI:** 10.64898/2026.06.30.735564

**Authors:** Nick Todd, Bridget Funk, Paige Nowlin, Christina Hung, Olaf Bodamer

## Abstract

Efficient delivery of molecular therapies to the central nervous system (CNS) remains a major barrier to treating neurogenetic disorders such as Niemann–Pick type C (NPC) disease. Focused ultrasound–mediated blood–brain barrier opening (FUS-BBBO) has emerged as a non-invasive strategy to enhance delivery of systemically administered therapeutics. In this study, we evaluated whether FUS-BBBO could enable delivery of lipid nanoparticle (LNP)-packaged modified mRNA (modRNA) to the cerebellum in an NPC mouse model.

A pilot study in wild-type mice demonstrated successful FUS-mediated BBB opening, delivery of LNP-packaged GFP mRNA, and subsequent protein expression in the cerebellum. We then performed a controlled study in NPC mice comparing delivery of LNP-GFP and LNP-NPC modRNA using intravenous administration with and without FUS-BBBO. BBB opening was confirmed by contrast-enhanced MRI in FUS-treated animals. Quantitative PCR revealed the presence of GFP mRNA in the cerebellum following FUS-BBBO, whereas NPC mRNA was minimal or undetectable across groups. However, no GFP or NPC1 protein expression was detected in the cerebellum by western blot in any experimental group. Consistent with this, no therapeutic effect on Purkinje cell survival was observed.

These results demonstrate that while FUS-BBBO reliably induces BBB opening and can facilitate limited delivery of LNP-packaged mRNA to the brain, this did not translate into detectable protein expression or therapeutic benefit in the NPC model under the conditions tested. This discrepancy between successful delivery in wild-type mice and lack of efficacy in diseased animals points to potential important biological and/or formulation-dependent barriers that must be addressed to enable effective CNS delivery of LNP-based mRNA therapies.

## Introduction

Niemann–Pick type C (NPC) disease is a rare, autosomal recessive lysosomal storage disorder caused predominantly by mutations in the *NPC1* gene, leading to impaired intracellular cholesterol trafficking and accumulation of lipids in multiple tissues, including the brain^1^. While visceral manifestations can be partially managed, progressive neurodegeneration—characterized by cerebellar ataxia, Purkinje cell loss, and cognitive decline—remains the primary driver of morbidity and mortality^2^. A central challenge in developing effective therapies for NPC and other neurogenetic disorders is the inability to deliver therapeutic agents across the blood–brain barrier (BBB), which restricts nearly all large molecules from entering the central nervous system.

Modified messenger RNA (modRNA) therapies have emerged as a promising approach for treating monogenic diseases by enabling transient expression of functional proteins without the risks associated with viral gene delivery^3–5^. For NPC, lipid nanoparticle (LNP)-packaged modRNA encoding NPC1 has been shown to restore protein expression and improve cellular function in vitro^6^ and in peripheral tissues in vivo. However, like many nanoparticle-based therapies, these formulations do not effectively cross the intact BBB, limiting their therapeutic impact on CNS pathology.

Focused ultrasound–mediated BBB opening (FUS-BBBO) is a non-invasive technique that uses ultrasound in combination with circulating microbubbles to transiently and locally increase BBB permeability^7–9^. This approach has been shown in preclinical^10–17^ and clinical studies^18–22^ to enable delivery of a wide range of therapeutics—including antibodies, nanoparticles, and viral vectors—to targeted brain regions. Importantly, FUS-BBBO offers the potential to combine systemic administration with spatially selective delivery, addressing key limitations of both invasive intracranial injections and diffuse systemic approaches.

In this study, we investigated whether FUS-BBBO could enable delivery of LNP-packaged modRNA to the cerebellum in a mouse model of NPC disease. Building on pilot data demonstrating successful delivery and protein expression of LNP-packaged GFP mRNA in wild-type mice, we performed a controlled study in NPC model mice to assess BBB opening, mRNA delivery, protein expression, and downstream therapeutic effects. By comparing intravenous administration with and without FUS-BBBO, this work aims to evaluate the feasibility of combining FUS-mediated delivery with modRNA therapeutics for treatment of CNS manifestations in NPC.

## Methods

### Mice and Experimental Design

All animal procedures were approved by the Brigham and Women’s Hospital Institutional Animal Care and Use Committee (IACUC) and conducted in accordance with institutional guidelines.

Two cohorts of mice were used. A pilot study was performed in wild-type BALB/c mice (N=11) to assess feasibility of focused ultrasound (FUS)-mediated delivery of lipid nanoparticle (LNP)-packaged mRNA. For the main study, NPC model mice (BALB/cNctr-Npc1m1N/J) were used to evaluate delivery and therapeutic effects. All mice underwent two treatment sessions, two weeks apart, and were sacrificed two days after the second treatment.

The full study was originally designed as a 3×2 factorial experiment with three routes of administration (intravenous (IV), IV combined with FUS-mediated blood–brain barrier opening (FUS + IV), and cisterna magna (CM) injection) and two LNP formulations (GFP and NPC1 modRNA). Group sizes were N=6 mice per group, which were completed for the two FUS + IV and IV only groups. However, CM injection proved technically challenging and associated with high mortality; only six mice underwent CM injection, and these data were excluded from analysis due to insufficient sample size and inconsistent delivery. Results for the main study are therefore presented over the 2 × 2 design of +/-FUS and LNP-GFP vs LNP-NPC. Outcome measures included BBB opening (MRI), mRNA delivery (qPCR), protein expression (western blot), and therapeutic effect (Purkinje cell counts) (**Figure 1**).

**Figure 1.**
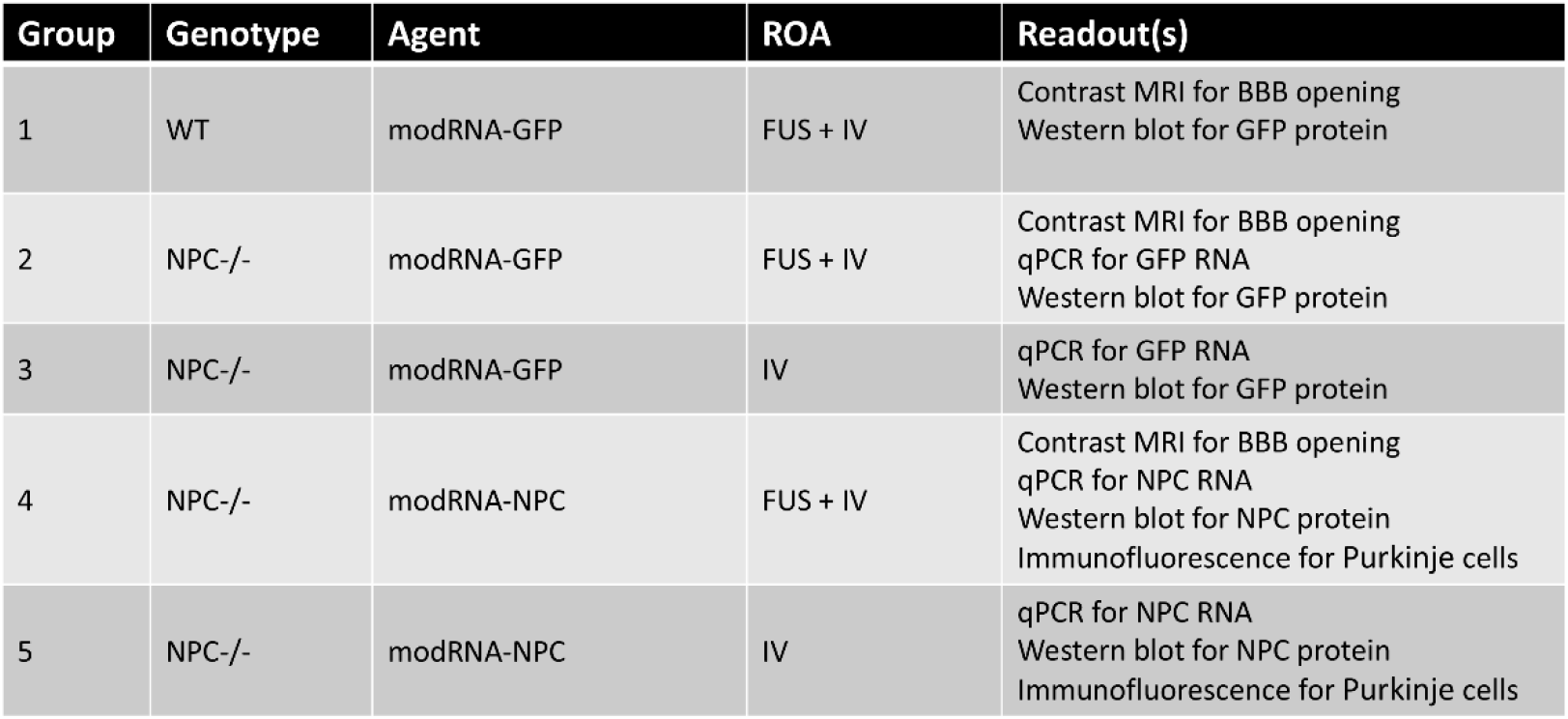
Experimental groups for the pilot study and the main study.

### FUS-BBB Opening and LNP Delivery

FUS-mediated BBB opening was performed using an in-house small-animal ultrasound system consisting of a single-element focused transducer operating at approximately 690 kHz, a positioning system, and a passive cavitation detector. Mice were anesthetized, and the head was immobilized in a stereotactic frame. Microbubbles (Optison, 100 µL/kg) were administered intravenously immediately prior to sonication. Ultrasound was applied using 10 ms bursts at 1 Hz repetition frequency for 120 seconds at an estimated peak negative pressure of ∼0.3 MPa. Four focal targets arranged in a grid pattern were applied to cover the cerebellum. Immediately following BBB opening, LNP formulations were administered via retro-orbital intravenous injection. In pilot experiments, LNP injection occurred immediately after sonication; this timing was maintained for the main study.

### LNP Formulations (GFP and modRNA-NPC1)

Two LNP formulations were evaluated: (1) LNP-packaged GFP mRNA as a reporter construct and (2) LNP-packaged modified mRNA encoding human NPC1 (modRNA-NPC1). The LNPs were provided by an external collaborator and based on previously characterized formulations optimized for mRNA delivery^23,24^. Particle size was approximately 80–100 nm^23^, consistent with prior reports of nanoparticles capable of crossing the BBB following FUS-mediated disruption. LNPs were administered at a dose of 1.5 mg/kg, within the range shown to be non-toxic in rats and non-human primates^23^.

### Read outs

#### Magnetic resonance imaging (MRI)

was used to confirm BBB opening in FUS-treated animals. T1-weighted contrast-enhanced images were acquired before and after intravenous administration of gadolinium contrast agent. Images were processed as percent signal change between pre- and post-contrast conditions. Regions of hyperintensity in the cerebellum were interpreted as evidence of BBB disruption

#### Quantitative PCR (qPCR)

was used to assess mRNA delivery to brain tissue. RNA was extracted from cerebellar and spleen samples, and expression of GFP and NPC1 transcripts was quantified relative to the housekeeping gene GAPDH and normalized to endogenous expression of NPC1 in a wild type mouse. Signal intensity thresholds were used to determine whether mRNA levels were above background. Positive and negative controls included spleen tissue and untreated brain samples.

#### Western Blot

was used to evaluate protein expression in cerebellar tissue harvested 48 hours after treatment. Samples were probed for GFP and NPC1 protein, with GAPDH used as the loading control. Protein expression in experimental groups was compared to a wild-type control to assess successful translation of delivered mRNA.

#### Immunohistochemical

analysis was performed on cerebellar sections to assess cellular outcomes. Purkinje cells were identified and quantified using stained sections, with cell counts normalized to tissue area. Representative sections were imaged, and Purkinje cell density was compared across experimental groups to evaluate potential therapeutic effects of NPC1 modRNA delivery.

## Results

### Pilot Study: FUS-BBBO Enables Delivery and Protein Expression of LNP-GFP in Wild-Type Mice

We first performed a pilot study in wild-type mice to assess whether FUS-mediated BBB opening could enable delivery of LNP-packaged mRNA to the cerebellum. Contrast-enhanced MRI demonstrated robust and spatially localized BBB opening in the cerebellum following FUS treatment, with consistent hyperintense signal observed across all treated animals, confirming successful BBB disruption (**Figure 2**). Western blot analysis of cerebellar tissue harvested 48 hours post-treatment revealed clear GFP protein expression in mice that underwent FUS-BBBO combined with LNP-GFP administration, whereas mice receiving LNP-GFP alone showed little to no detectable signal (**Figure 3**). These results demonstrate that FUS-BBBO enables both delivery of LNP-packaged mRNA across the BBB and subsequent translation into protein in the targeted brain region.

**Figure 2.**
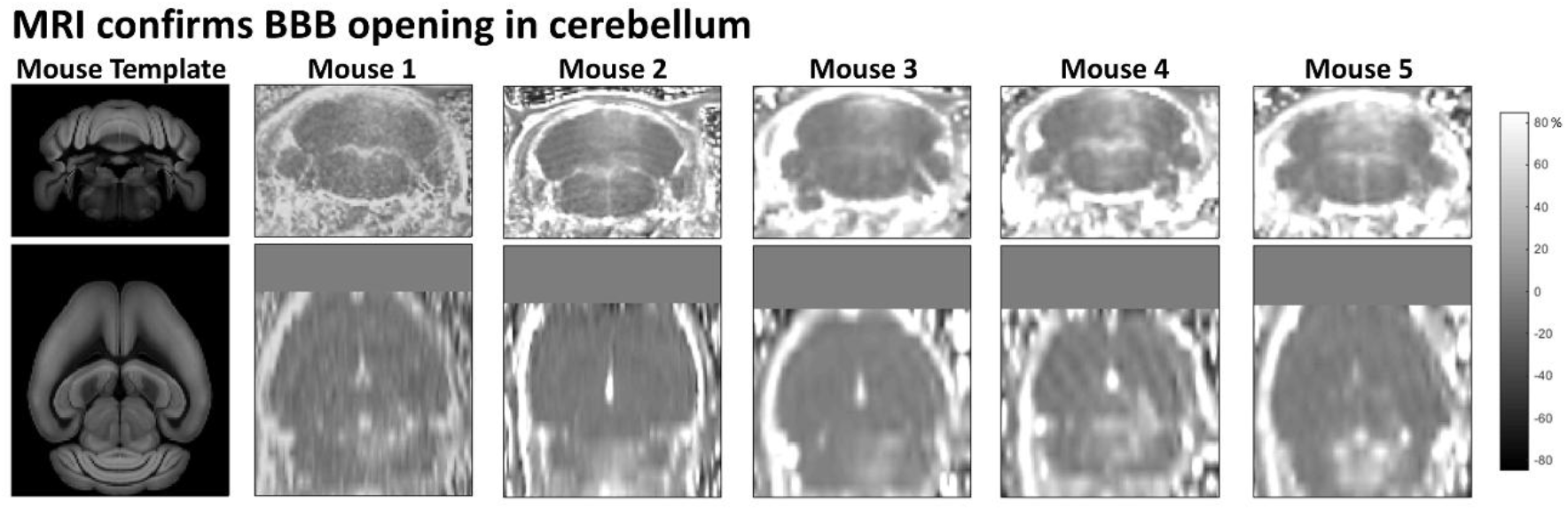
T1-weighted contrast MRI for mice in the pilot study that underwent FUS-BBB opening treatment, confirming BBB opening in the cerebellum.

**Figure 3.**
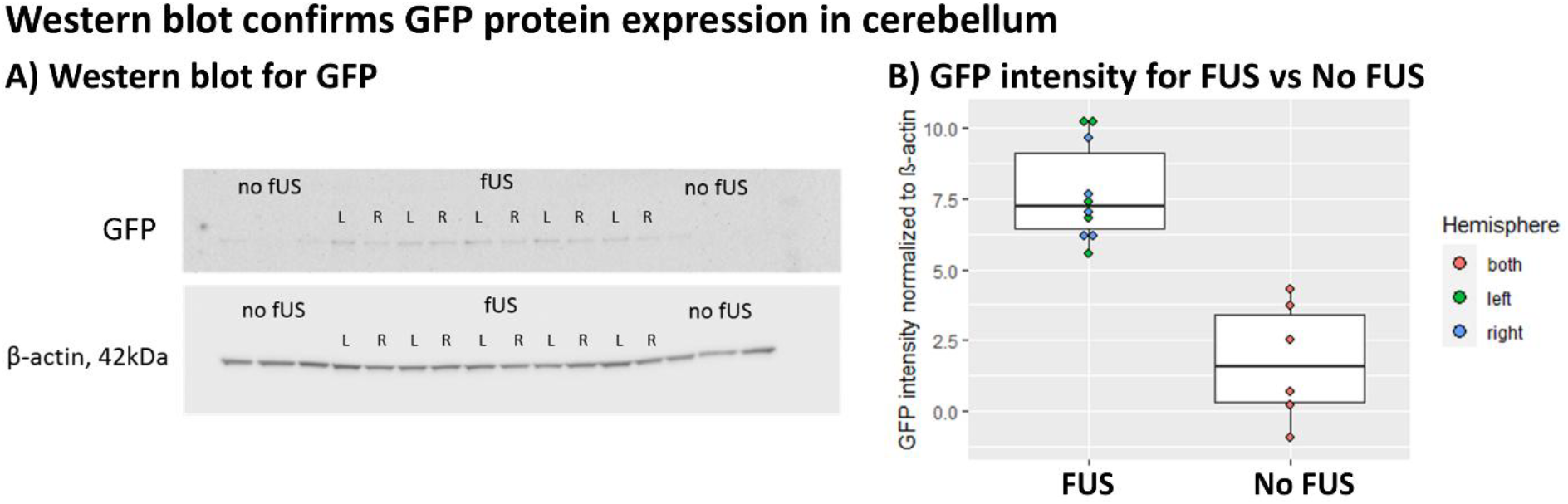
Western blot results for the pilot study assessing GFP expression in cerebellum. GFP signal, normalized to β-actin, was significantly higher in the FUS-treated group compared to mice who did not receive FUS.

**Figure 4.**
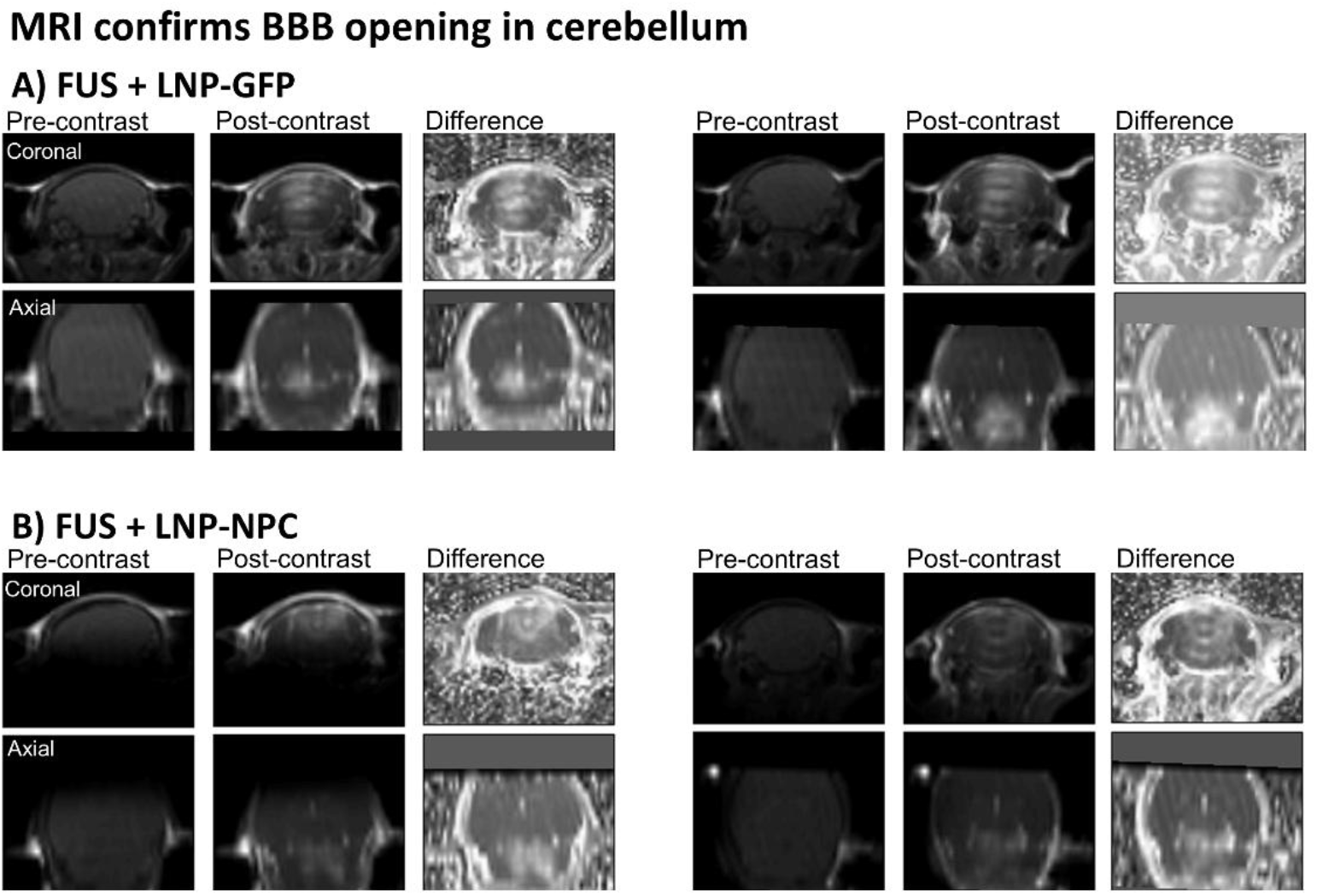
Example MRI images from two mice each in the FUS + LNP-GFP and FUS + LNP-NPC groups. Images show pre-contrast T1-weighted MRI, post-contrast T1-weighted MRI and the difference. Hyperintense signal in the cerebellum confirms BBB opening.

### Full Study in NPC Mice

In the full study, we evaluated FUS-mediated delivery in NPC model mice. Contrast-enhanced MRI confirmed successful BBB opening in the cerebellum of FUS-treated animals of both FUS + LNP-GFP and FUS + LNP-NPC cohorts. All imaged mice showed localized increases in signal intensity in the cerebellum following gadolinium administration, indicating consistent BBB disruption across the cohort.

To assess delivery of LNP-packaged mRNA, we quantified GFP and NPC1 transcripts by qPCR in cerebellar and spleen tissues collected from all treatment groups. GFP and NPC1 probe sets were applied to each sample, generating six measurements for each probe: cerebellum from GFP-IV, GFP-IV+FUS, NPC-IV, and NPC-IV+FUS groups, as well as spleen samples from GFP- and NPC-treated animals (Figure 5A). This design provided both positive and negative controls for each assay. For example, spleen tissue from GFP-treated mice served as a positive control for GFP detection and a negative control for NPC1 detection, whereas spleen tissue from NPC-treated mice served as a positive control for NPC1 detection and a negative control for GFP detection.

**Figure 5.**
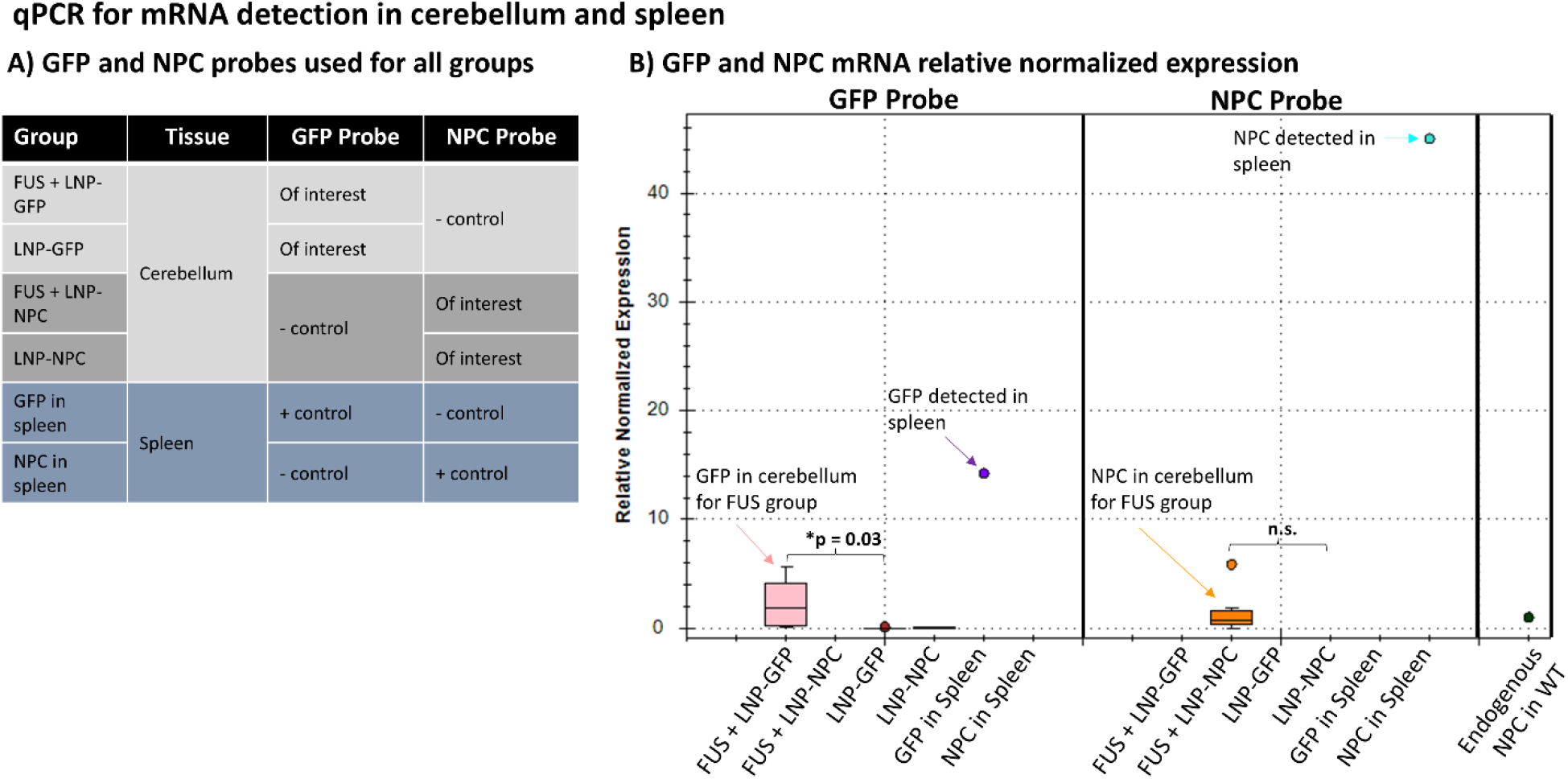

Transcript levels were normalized to GAPDH and expressed relative to endogenous Npc1 expression measured in wild-type mouse cerebellum (Figure 5B). As expected, strong GFP signal was detected in spleen tissues from GFP-treated animals and strong NPC1 signal was detected in spleen tissues from NPC-treated animals, confirming probe specificity and successful detection of exogenous transcripts. In mice receiving LNP-GFP with FUS-BBBO, GFP mRNA was detected in the cerebellum at levels exceeding those observed in non-FUS controls (p=0.03), indicating successful FUS-mediated delivery of GFP mRNA to the targeted brain region.

For the therapeutic construct, NPC1 mRNA was readily detected in spleen samples from NPC-treated animals, confirming successful systemic delivery and assay performance. However, NPC1 mRNA levels in the cerebellum were low to undetectable in both IV and IV+FUS groups, with no significant difference between groups. These findings indicate that while FUS-BBBO enabled detectable delivery of GFP mRNA to the brain, delivery of NPC1 mRNA was substantially less efficient under the same experimental conditions.

Despite evidence of mRNA delivery in some conditions, western blot analysis revealed no detectable GFP or NPC1 protein expression in the cerebellum of NPC mice across all experimental groups (**Figure 6**). In contrast to the pilot study in wild-type mice, neither reporter nor therapeutic protein was observed following FUS-BBBO-mediated delivery. NPC1 protein was detected in wild-type control tissue, as expected, but was absent in NPC model mice regardless of treatment condition. These results suggest that delivered mRNA did not result in sufficient protein translation in the diseased brain under the conditions tested.

**Figure 6.**
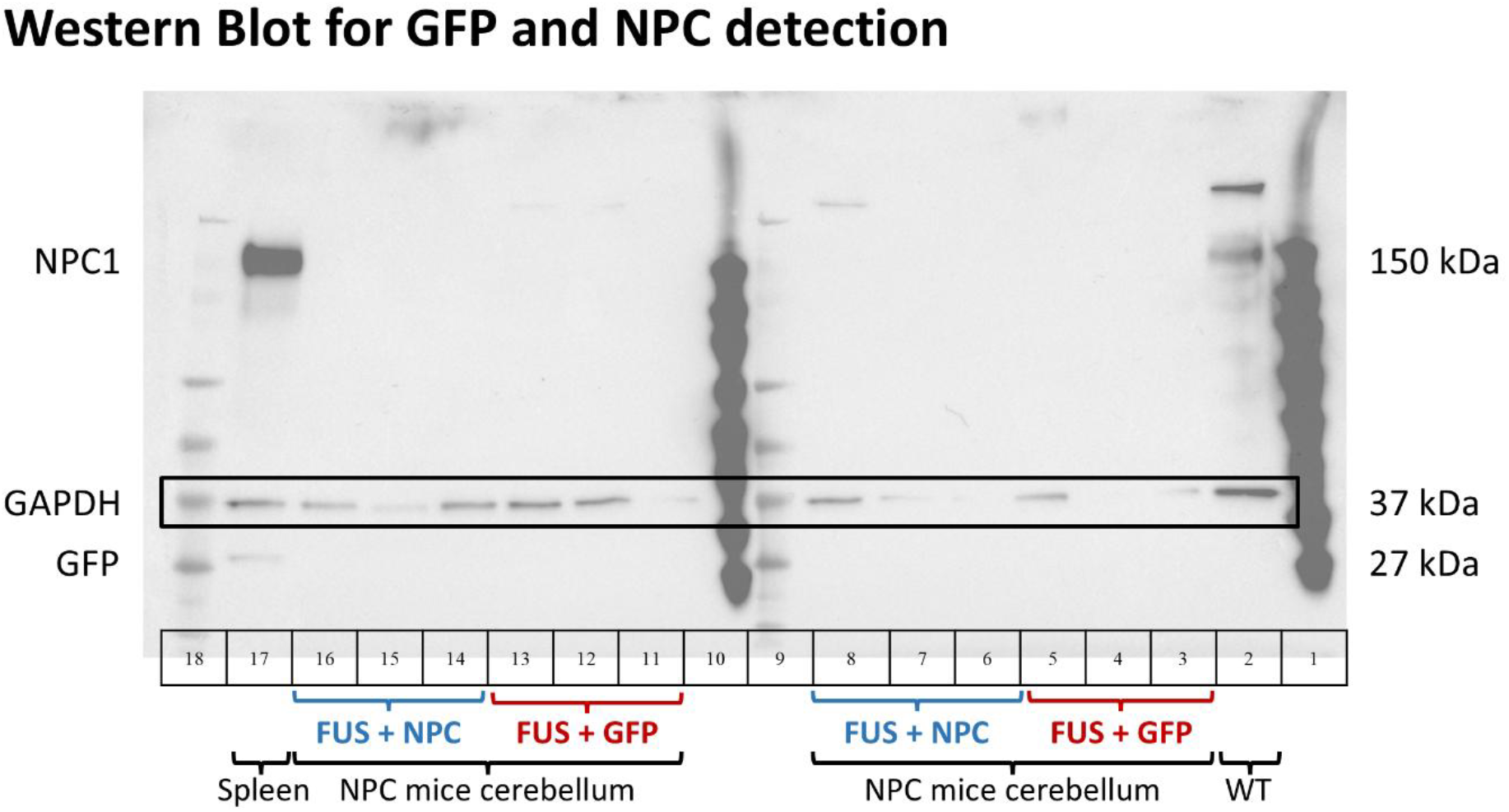
Western blots for FUS + LNP-GFP and FUS + LNP-NPC groups. NPC signal at ∼150 kDa is seen in two control conditions: spleen of an NPC mouse (#17) and cerebellum of a wild-type mouse (#2). NPC signal is not seen in the cerebella of any NPC mice. GFP signal @ 27 kDa was not seen in the FUS-treated NPC mice.

To assess functional impact of therapy, we quantified Purkinje cell counts in the cerebellum using immunohistochemistry (**Figure 7**). No significant differences in Purkinje cell counts were observed between treatment groups, including mice receiving LNP-NPC with FUS-BBBO. Consistent with the absence of detectable NPC1 protein expression, these findings indicate that the delivered therapy did not produce a measurable therapeutic effect on cerebellar pathology in this model.

**Figure 7.**
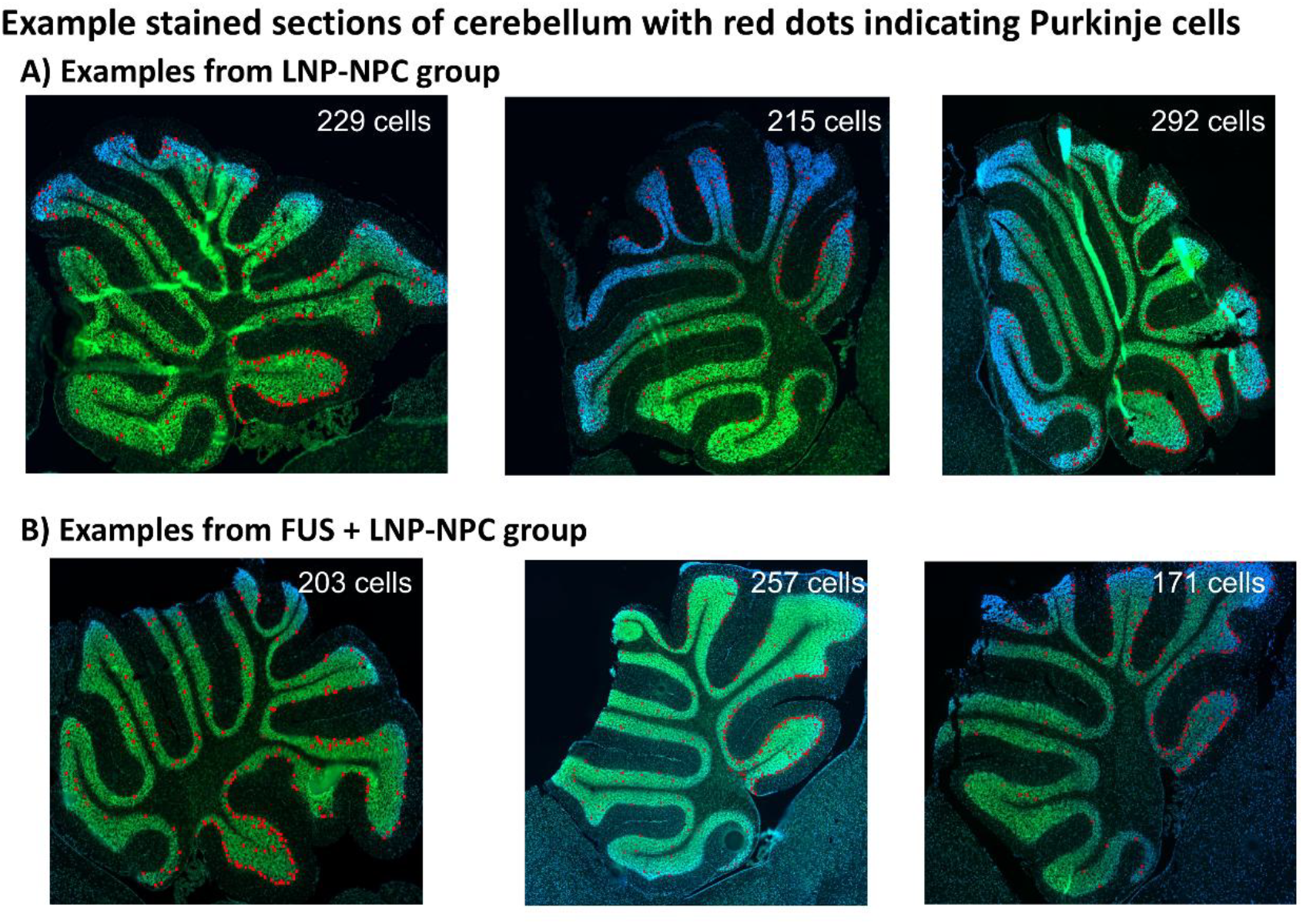
Purkinje cell death is one of the most prominent manifestations of the neuropathology of NPC seen in mice. These example images show cell counts of purkinjie cells in cerebellum labeled with red dots.

## Discussion

In this study, we evaluated whether focused ultrasound–mediated blood–brain barrier opening (FUS-BBBO) could enable delivery of lipid nanoparticle (LNP)-packaged modified mRNA to the cerebellum in a mouse model of Niemann–Pick type C (NPC) disease. Our results demonstrate a mixed outcome. In wild-type mice, FUS-BBBO enabled delivery and translation of LNP-packaged GFP mRNA in the cerebellum. In contrast, in NPC model mice, although BBB opening was robust and reproducible based on contrast MRI, limited mRNA delivery could be detected and this did not translate into detectable protein expression or therapeutic benefit. Together, these findings highlight both the promise and the current limitations of combining FUS-BBBO with LNP-based nucleic acid therapies for central nervous system (CNS) delivery.

A key strength of this study is the clear demonstration that FUS-BBBO remains a reliable and consistent method for transiently increasing BBB permeability, even in a diseased brain and difficult to reach region such as the cerebellum. MRI-confirmed BBB opening was observed in all treated animals, consistent with a large body of preclinical and clinical literature showing that FUS with microbubbles can safely and reproducibly disrupt the BBB in a spatially targeted manner. This reinforces the role of FUS as a robust physical delivery modality that can, in principle, enable entry of otherwise impermeable therapeutics into the brain.

However, our findings also underscore that BBB opening alone is not sufficient for effective therapeutic delivery, particularly for nanoparticle-based nucleic acid therapies. While GFP mRNA was detectable in the cerebellum following FUS-BBBO, delivery of NPC1 mRNA was minimal, and neither construct resulted in detectable protein expression in NPC mice. This disconnect between BBB opening, molecular delivery, and functional protein expression suggests both the need for further optimization of the combined LNP vector and FUS-BBB opening protocol and the possible presence of additional biological and/or formulation-dependent barriers beyond vascular permeability.

These observations are consistent with prior work demonstrating that successful FUS-mediated delivery of nanoparticles depends critically on nanoparticle design. For example, recent studies have shown that long-circulating, serum-stable nanoparticles with optimized surface properties (e.g., dense PEGylation) can accumulate in FUS-targeted brain regions and achieve robust nucleic acid expression, whereas clinically relevant lipid nanoparticles may fail to do so despite equivalent BBB opening. In particular, Kwak et al. demonstrated that biodegradable PEG–PBAE nanoparticles enabled efficient mRNA and DNA delivery with subsequent protein expression in neurons and astrocytes, while LNP formulations analogous to those used clinically showed limited brain delivery under similar conditions^25^. These findings suggest that circulation time, serum stability, and interactions with the brain extracellular matrix are key determinants of delivery success.

More broadly, prior studies combining FUS with polymeric or lipid-based nanoparticles have demonstrated that particle size, surface chemistry, and diffusivity within the brain parenchyma critically influence therapeutic distribution and efficacy. Nanoparticles on the order of ∼40–100 nm with dense PEG coatings have been shown to penetrate brain tissue effectively following FUS-BBBO, whereas larger or more adhesive particles exhibit limited distribution^26^. In addition, even after crossing the BBB, nanoparticles must navigate the dense extracellular matrix of the brain, which can significantly restrict diffusion and cellular uptake.

The discrepancy between the pilot study in wild-type mice and the lack of protein expression in NPC mice further suggests that disease-specific factors may play a critical role in limiting therapeutic efficacy. NPC pathology is associated with widespread lipid accumulation, altered endosomal–lysosomal trafficking, and neuroinflammation, all of which could impair nanoparticle uptake, intracellular trafficking, or mRNA translation. It is also possible that differences in vascular integrity, immune activation, or cellular metabolism in the diseased brain influence both delivery and expression of LNP-packaged mRNA. These factors may contribute to the observed failure to achieve detectable protein expression despite evidence of mRNA presence.

Several additional factors may have contributed to the limited efficacy observed in this study. First, the LNP formulation used may not have been optimized for brain delivery following systemic administration and FUS-BBBO, particularly with respect to circulation half-life and resistance to serum degradation. Second, the timing and dosing of LNP administration relative to BBB opening may not have maximized delivery efficiency. Third, BBB opening, while detectable in all animals, may not have sufficiently covered the full cerebellum and thus diluted the signal of mRNA and protein expression. Fourth, the level of mRNA delivered to the cerebellum may have been below the threshold required for detectable protein expression or therapeutic effect. Finally, technical challenges associated with alternative delivery routes, such as cisterna magna injection, limited the ability to fully evaluate complementary strategies.

FUS-BBBO provides a powerful and reliable method for enabling access to the brain, but successful therapeutic delivery of nucleic acids requires careful integration of delivery physics with nanoparticle engineering and disease biology. Future work should focus on optimizing nanoparticle formulations for prolonged circulation and enhanced brain penetration, improving intracellular delivery and translation efficiency, and understanding how disease-specific factors influence therapeutic response. Combining FUS-BBBO with next-generation nanoparticle platforms will likely enable effective, non-invasive delivery of gene therapies to the brain.

## Acknowledgements

This study was supported by the Focused Ultrasound Foundation.

